# Predicting neuropsychiatric symptoms of Parkinson’s disease with measures of striatal dopaminergic deficiency

**DOI:** 10.1101/763110

**Authors:** Ram Bishnoi, Marina C. Badir, Sandarsh Surya, Nagy A. Youssef

## Abstract

The role of nigrostriatal dopaminergic neurons degeneration is well established in the pathophysiology of Parkinson’s disease. However, it is unclear if and how the degeneration of the dopamine pathways affects the manifestation of the neuropsychiatric symptoms (NPS) of Parkinson’s disease (PD). Dopamine transporter (DAT) imaging, a technique to measure the reduction in the dopamine transporters is increasingly used as a tool in the diagnosis of PD. In this study, we examine if the baseline dopamine transporter density in the striatum measured by striatal binding ratio (SBR) is associated with the longitudinal onset and/or progression of NPS in PD as measured by the part 1 of Movement Disorder Society - Unified Parkinson’s Disease Rating Scale, over four years. Data of patients with PD and an abnormal screening present on 123I-ioflupane single-proton emission computed tomography were obtained from Parkinson’s Progression Markers Initiative (PPMI) database. Latent Growth Modeling (LGM), a statistical technique that can model the change over time while considering the variability in rate of change at the individual level was used to examine the progression of NPS over time. The results indicate the SBR did not correlate with the baseline NPS but did correlate with the rate of change of NPS (p<0.001) over the next four years, even after eliminating age related variance which can be a significant confounding factor. In conclusion, this study showed gradual worsening in NPS in patients with Parkinson’s disease which inversely correlates with the density of the dopamine transporters as measured by SBR at baseline.

## INTRODUCTION

Parkinson’s Disease (PD) is typically characterized by motor symptoms, however neuropsychiatric symptoms (NPS) are not uncommon in individuals with PD. NPS have significant impact on individuals’ quality of life and caregiver burden (1). Commonly observed NPS in PD are- depression, anxiety, apathy, hallucinations, cognitive impairments etc. These symptoms are present in up to 40% of individuals diagnosed with PD (2) and are integral part of PD pathophysiology. They are seen early in the course of illness, even before the onset of motor symptoms of PD (3). Presence of NPS has shown to predict future deterioration of cognitive and the motor symptoms (4, 5).

Neurodegenerative changes involving dopaminergic system are directly responsible for motor symptoms in PD (6). Visualization of striatal dopamine transporters (DAT) has been extensively studied as potential evaluation tool in PD; to establish nigrostriatal dopaminergic degeneration in patients displaying motor signs and symptoms of parkinsonism (7). The role of dopaminergic system in NPS in PD is not well understood. Previous studies have shown that alterations in striatal uptake of the DAT may have association with NPS (8–11), however the results are mixed in this context (11–13). Most of these studies have examined the correlation between NPS and DAT imaging, cross-sectionally. In this study, we aimed to investigate the impact of dopamine striatal binding on the longitudinal course of NPS in PD over four years. We hypothesized that NPS gradually worsen during the course of PD and dopaminergic depletion as measured by DAT scan play an important role in their onset and progression.

## METHODS

### Study population

Data used for analysis in this study were obtained from the PPMI database (for up‐to‐date information on the study, visit www.ppmi-info.org). The PPMI is an international, multicenter, prospective study designed to discover and validate biomarkers of disease progression in newly diagnosed PD participants (National Clinical Trials identifier NCT01141023). Each PPMI recruitment site received approval from an institutional review board or ethics committee on human experimentation before study initiation. Written informed consent for research was obtained from all individuals participating in the study. Inclusion criteria for the PPMI consisted of a clinical diagnosis of PD, age older than 30 years, Hoehn and Yahr Stage 1 or 2, diagnosis within 24 months, and an abnormal ^123^I‐ioflupane single‐proton emission computed tomography scan at screening.

### Assessments

#### Neuropsychiatric symptoms

A commonly used instrument in quantitative phenotyping of PD is the Movement Disorder Society - Unified Parkinson’s Disease Rating Scale (MDS-UPDRS), which has 4 subscales - Part I, Non‐Motor Aspects of Experiences of Daily Living (13 items); Part II, Motor Aspects of Experiences of Daily Living (13 items); Part III, Motor Examination (33 items); and Part IV, Motor Complications (6 items) (14).

Part I of MDS-UPDRS assesses non-motor symptoms. Part IA is rater-administered and consists of following 6 questions focusing on complex behaviors: Cognitive impairment, Hallucinations and psychosis, Depressed mood, Anxious mood, Apathy, and features of dopamine dysregulation syndrome. All six questions are rated on a scale of 0 to 4 based on severity of symptoms. This sub-scale includes commonly observed neuropsychiatric symptoms of Parkinson’s disease and we considered total score from 6 questions as measure of NPS in our analysis.

### Dopamine transporter imaging

DAT SPECT imaging (DatScan) was acquired and analyzed from all subjects at baseline as per PPMI imaging protocol. Details of the protocol can be accessed at http://www.ppmi-info.org.

Raw SPECT data were subjected to reconstruction, attenuation correction and normalization to MNI space. Region of interests included right and left caudate, right and left putamen, and cerebellum (reference tissues). Count densities were extracted from each of those regions and used to calculate striatal binding ratio (SBR) for each of those four regions. We used mean of all four regions as SBR in our analysis.

### Statistical analysis

Descriptive statistics were analyzed using SPSS (version 25). To test whether the NPS progressed over time, Latent Growth Modeling (LGM) was used with the support of AMOS (version 26). LGM is a flexible analytic technique for modeling change over time, which takes variability in rate of change at the individual level into account and focuses on correlations over time, changes in variances and in mean values (15).

A major advantage is that LGM can handle missing data, as it uses data from all participants, not only from those who have completed all four years of assessments, and as such provides less biased information on prediction results(16).

Following guidelines for longitudinal assessment (Little 2013), model fit was evaluated based on comparative fit index (CFI), and the root mean square error of approximation (RMSEA). Acceptable fit to the data was indicated by CFI values ≥ 0.90 and RMSEA values ≤ 0.08. (17, 18).

## RESULTS

Demographic and participants characteristics are shown in Table 1. The base univariate model analysis (model 1) is shown in figure 1. The intercept represented mean score of NPS at baseline. The slope was assumed to have linear growth, as the loading slope factors were fixed at values of 0, .25, .5 and 1 representing assessment at baseline, year 1, year 2 & year 4, respectively. The model fit of base model was satisfactory, and it was determined that the slope factor for NPS estimated linear change. The univariate linear growth models show that NPS had an mean initial score (intercept) of 1.27 (*p* < .001), with an increase (slope) of 1.02 (*p* < .001). Model 2 presents the results of linear growth curve model with baseline mean striatal SBR as a predictor of initial score and rate of change of NPS. Path diagram of this model is shown in figure 2. The results indicate a significant impact of striatal SBR on rate of change of NPS (p<0.001) but not the mean baseline score.

**Table 1.** Demographic and assessment variables of participants.

|  |  |
| --- | --- |
| Parkinson's disease, N | 423 |
| Age, Mean (SD) | 61.6 (9.7) |
| Sex, Male N(%); Female N(%) | 277 (65): 146 (35) |
| Education, Mean number of years (SD) | 15.56(2.97) |
| MDS-UPDRS, Part-I |  |
| Baseline, N (missing data) | 423 (0) |
| Year 1, N (missing data) | 367 (56) |
| Year 2, N (missing data) | 365 (58) |
| Year 4, N (missing data) | 342 (81) |
| Mean striatal specific binding ratio (SBR) |  |
| Missing data | 4 |
| Mean (SD) | 1.41(.40) |

**Fig 1.**
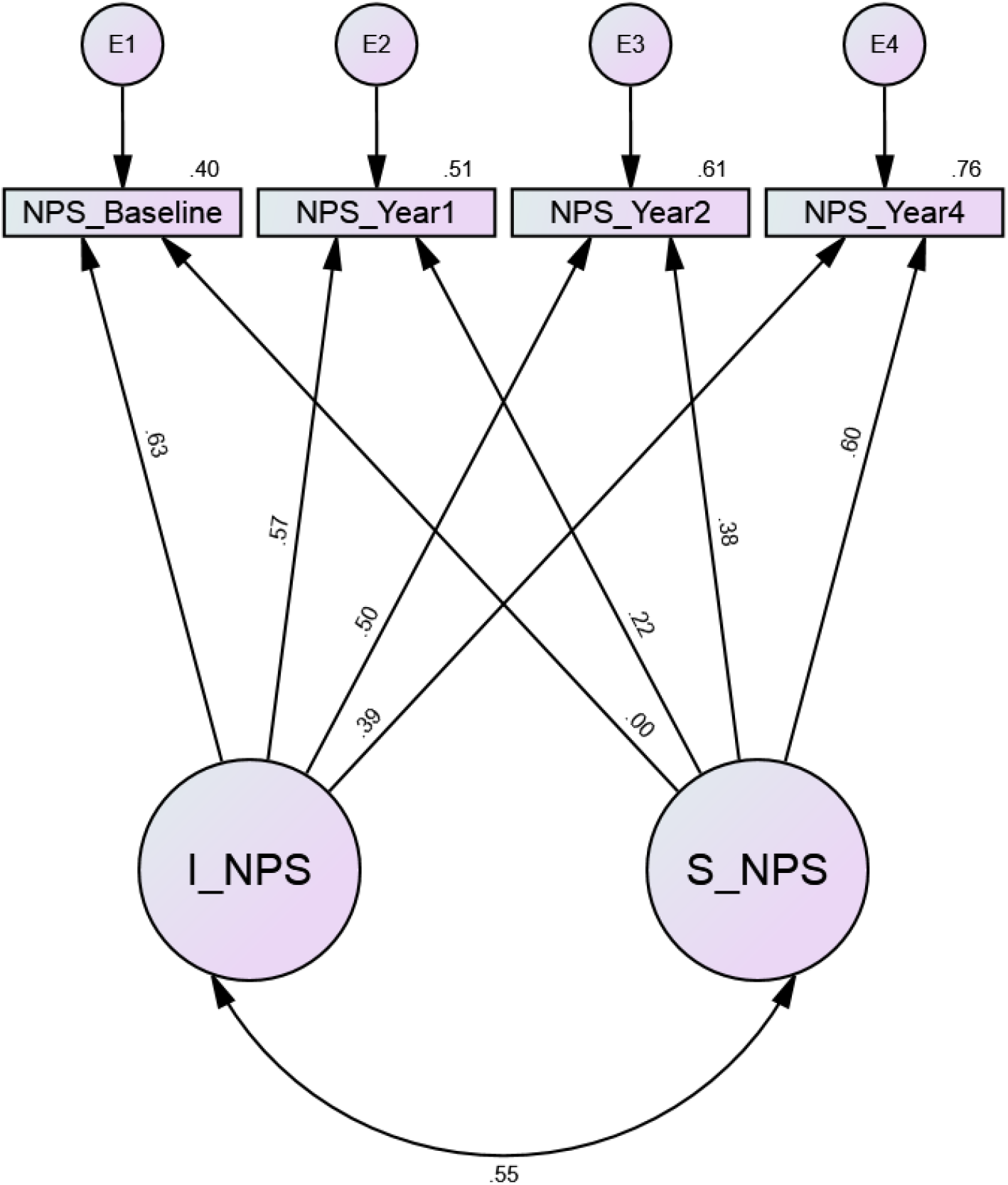
Model 1. Latent growth model exploring rate of change of neuropsychiatric symptoms (NPS) in Parkinson’s disease (PD). I_NPS: intercept of NPS; S_NPS: Slope of intercept. Model fit: χ^2^(8)= 27.42, CFI= 0.96, RMSEA= 0.07; \**p* < 0.05, \*\**p* < 0.01, \*\*\**p* < 0.001; standardized path coefficients are presented.

**Fig 2.**
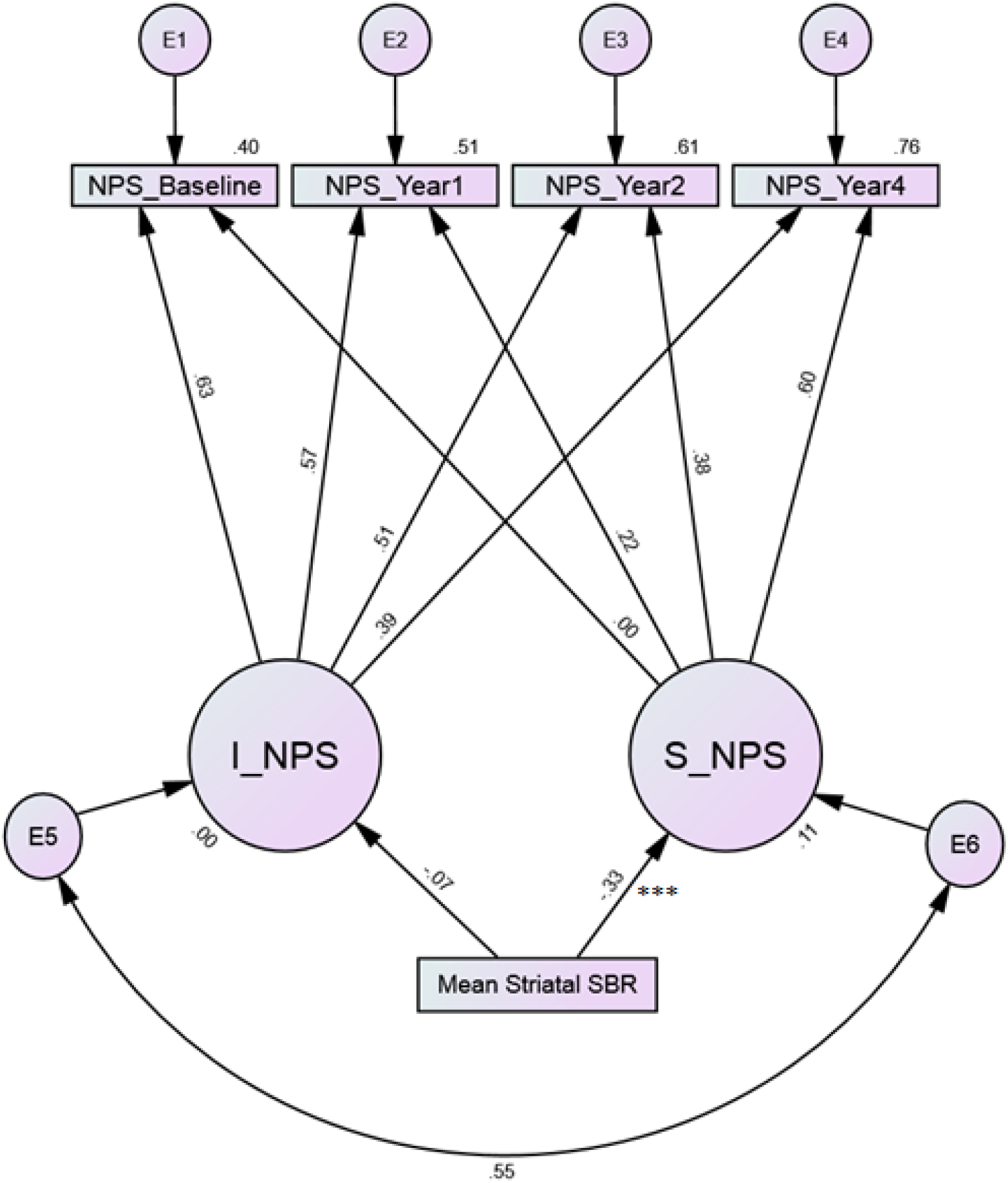
Model 2. Conditional latent growth model with mean striatal SBR as an exogenous predictor. I_NPS: intercept of neuropsychiatric symptoms (NPS); S_NPS: Slope of intercept. Model fit: χ^2^(10)= 28.70, CFI= 0.96, RMSEA= 0.07; \**p* < 0.05, \*\**p* < 0.01, \*\*\**p* < 0.001; standardized path coefficients are presented.

Age is known to have its independent effect on several neuropsychiatric symptoms included in analysis here including cognition, depression, anxiety etc. (19). In model 3, as shown in figure 3, we used age as an additional predictor of NPS’ intercept and slope. In this patient population with mean age of 61.6 years, age was not a significant predictor of baseline NPS score (p=0.83) or rate of change of NPS (p=0.05). Mean striatal SBR significantly predicted slope (p<0.001) even after eliminating age related variance from NPS scores.

**Fig 3.**
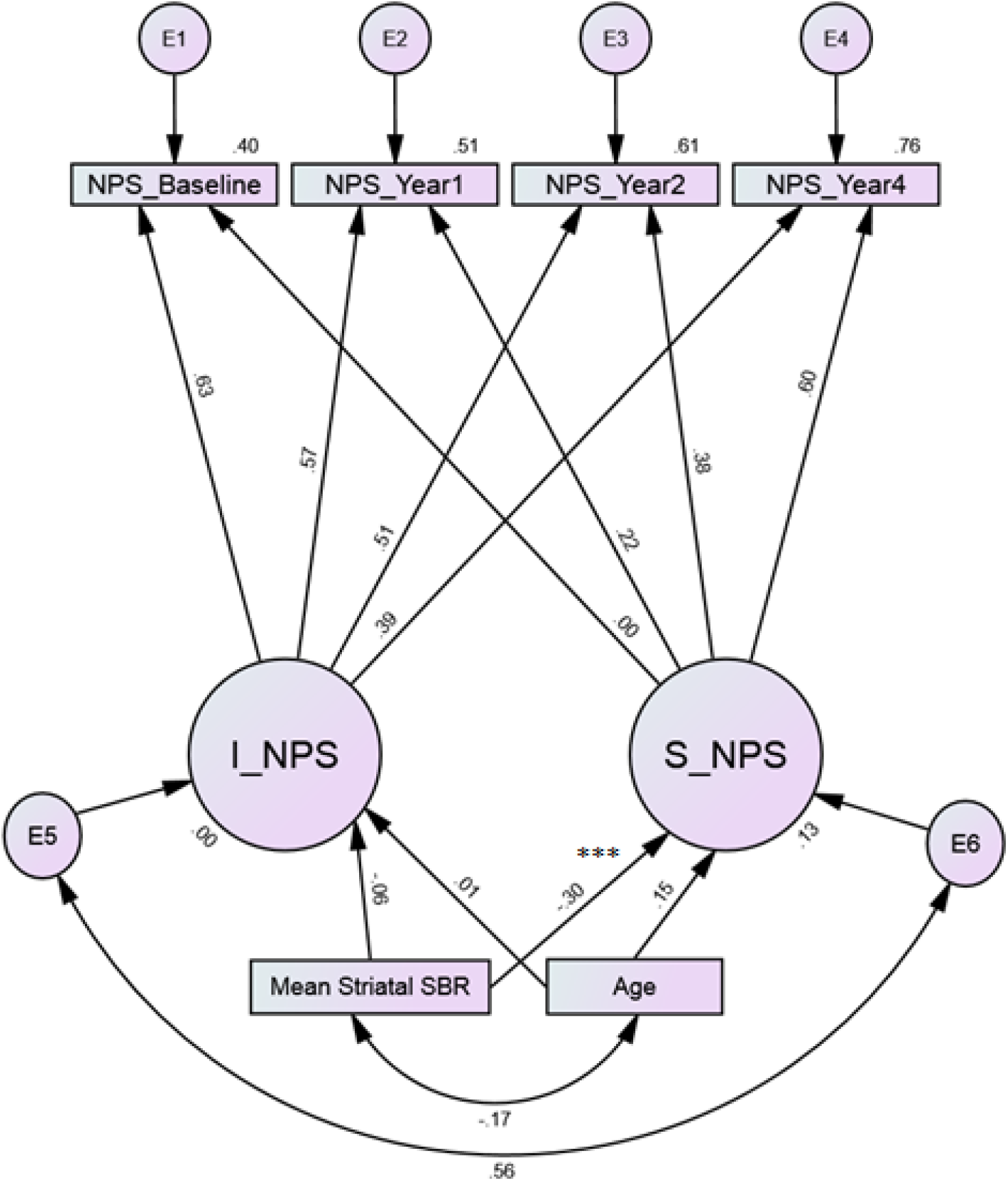
Model 3. Conditional latent growth model with mean striatal SBR and age as exogenous predictors. I_NPS: intercept of neuropsychiatric symptoms (NPS); S_NPS: Slope of intercept. Model fit: χ^2^(12)= 35.55, CFI= 0.95, RMSEA= 0.06; \**p* < 0.05, \*\**p* < 0.01, \*\*\**p* < 0.001; standardized path coefficients are presented.

## DISCUSSION

We examined the relationship of striatal dopaminergic deficit with NPS over a four-years period in 423 PD subjects. Gradual worsening of NPS in patients with Parkinson’s disease is significant over time in the base model. Dopamine depletion as measured with DAT imaging, significantly impacted the longitudinal course of NPS in PD. Striatal DAT binding measured at baseline did not predict the NPS at baseline, but does predict the longitudinal changes in NPS when measured for four years. These results were significant after adjusting for age. Non-significant association of SBR with intercept of NPS is not a novel finding. Several previous studies have reported similar results (11, 12, 20) where NPS in untreated PD patients were not significantly associated with DAT scan results. However, most salient finding of our analysis is that the SBR at baseline significantly predict the future trajectory of NPS over 4-year period. Significant and negative association between SBR and NPS indicates that lower striatal DA availability leads to greater trajectory of NPS. Given that age is known to have its independent effect on several neuropsychiatric symptoms, age as additional predictor for NPS’ intercept and slope did not change the results.

Although there are multiple studies in the literature that examine Parkinson’s patients using DAT on cognitive outcomes, our literature search did not find any studies specifically examining NPS in patients with Parkinson’s disease using FDA approved DAT scans that were longitudinal in design and included a large number of patients. However, related studies have been done in an attempt to examine the predictive power of prognostic scans in patients with PD. A 2012 retrospective study by Ravina et al. aiming “to measure the prognostic value of imaging for motor and nonmotor outcomes in Parkinson’s disease (PD)” examined the long-term motor and nonmotor symptoms of 537 patients with early, untreated PD. The researchers correlated motor and nonmotor symptoms with baseline striatal binding of dopamine in each patient. They evaluated dopamine striatal binding using DAT imaging with (^123^I) (β)-CIT and single photon emission computerized tomography (SPECT). The nonmotor symptoms assessed included the mental status, cognitive ability, depressive symptoms, and quality of life. The results showed significant correlation between baseline striatal binding and both motor and psychiatric symptoms after 5 or 6 years. Low baseline striatal binding correlated with a higher likelihood of motor, depressive, and psychotic symptoms, and cognitive impairment (21). However, this was the initial exploratory study and will require future replications to confirm these finding; as the authors suggested that the results should be “treated as hypothesis generating and require confirmation”.

In regard to the NPS of anxiety and depression, Weintraub et al. found an inverse correlation between high anxiety and depression scores and low left anterior putamen DAT availability in patients with PD (22). The study compared the striatal DAT SPECT brain scans, State-Trait Anxiety Scale, and the Profile of Mood States (POMS) of 76 patients with PD and 46 healthy volunteers. The results of the study suggested that lower levels of striatal dopamine, particularly in the left anterior putamen, was inversely coordinated with more adverse NPS of anxiety and depression in patients with PD (22). The association between total affect scores and DAT availability was found to be present only in the subgroup of patients with less severe PD (r=-0.35; P=0.04). However, participants with the highest DAT availability did not have "high total affect scores". Although Weintraub et al. focus solely on anxiety and depression, the results support the importance of dopaminergic systems of the progression of NPS in patients with PD.

Another smaller retrospective study that consecutively reviewed PD patients who underwent both brain magnetic resonance imaging (MRI) and ^18^F-radiolabeled N-(3-fluoropropyl)-2β-carboxymethoxy-3β-(4-iodophenyl) nortropane (^18^F-FP-CIT) positron emission tomography (PET) to examine the relationship between dopamine depletion and “non-motor symptoms” in patients with PD failed to find an association between striatal dopamine depletion and “non-motor” symptoms of PD such as depression, anxiety, fatigue, sleep quality, global cognition, and frontal executive function (23). Unlike our study, which utilized DaTscan SPECT imaging technology, the Park et al. study evaluated striatal dopamine depletion through DAT imaging with PET ^18^F-FP-CIT, which allowed them to assess striatal dopamine in 12 sub-regions. Forty-one patients with PD were included in the analysis to determine whether there was a correlation between striatal dopamine depletion and non-motor symptoms. The study did not find correlation between striatal dopamine depletion and non-motor symptoms. It should be noted that the Park et al. study included patients who were taking dopaminergic medications at the time of the study, and the researchers reported that they “could not exclude a possibility of the effect of dopaminergic medication on various non-motor symptoms evaluated in the current study” (23). It should also be noted the Park et al. study was probably underpowered to detect differences even if they existed considering the sample size was only 41 patients and the study utilized multiple analysis across 12 striatal sub-regions.

Perhaps combining imaging biomarkers with clinical symptoms could enhance our predictive ability of the course of these symptoms and the trajectory of neuropsychiatric symptoms. For instance, a 2019 pilot study consisted of 262 PD patients from the PPMI database who showed no signs of cognitive impairment at baseline, had accessible [^123^I] FP-CIT SPECT and CSF data, and completed a 36-month follow-up to assess signs of cognitive decline. The study found that when CSF Aβ42, CSF total tau, and [^123^I]FP-CIT caudate binding) are used individually they are not as helpful, however, when combined the researchers reported that this “pilot study demonstrated that a combined evaluation of biological and neuroimaging markers enables the identification of 65% of drug-naïve PD patients that will develop CI over a 36-month follow-up.” (24).

Our study has several limitations as follows:

1. We did not look at individual neuropsychiatric disorders on this analysis thus the resulting heterogenicity may have changed the direction of association of a single disorder. However, given the role of dopamine in several of the neuropsychiatric disorders that develop during the course of PD we thought it would be helpful to establish first this relationship.
2. We only aimed to examine the correlation between dopamine depletion and progression of NPS of PD. Serotoninergic neurons modulate the dopamine release in cortical and subcortical areas of the brain (De Deurwaerdère and Di Giovanni 2017) and serotonergic degeneration do have a role in the motor symptoms and NPS of PD (Politis and Niccolini 2015). This study did not examine the effects of other neurotransmitter systems on the NPS of PD and unclear how examining these systems will impact the predictability of NPS in PD by dopamine depletion alone.
3. The rating scales used in this study measured disorders but not assess any RDoC constructs (25, 26) in a cross diagnostic fashion. Future, studies may explore such associations between RDoC constructs and PD. Again, we thought it would be necessary to first examine associations with well-established diagnostic entities supported by a vast literature.
4. There may have been confounder that were not controlled for in this non-experimental non-randomized design of the study and might have affected the results.

None the less one of the important strengths of this study is that it is the largest study in the in the literature to examine this question. In addition, the prospective longitudinal design with 8-year follow-up with 4 assessment using imaging is also a strength. Moreover, the imaging methodology used in the study has been FDA approved specifically for assisting PD diagnosis.

In conclusion, this study showed gradual worsening in NPS in patients with Parkinson’s disease. This study also shows that change in SBR significantly predicted the course of NPS. Thus, this study suggests that, in Parkinson’s disease, the dopaminergic system plays an important role in NPS and its course. Mean striatal SBR also significantly predicted change in NPS symptoms (even after eliminating age related variance). Future larger studies are needed to replicate these finding and to examine separate correlation with individual psychiatric disorders as well as RDoC constructs.

## Acknowledgment

PPMI - a public-private partnership - is funded by the Michael J. Fox Foundation for Parkinson’s Research and funding partners, including Abbvie, Avid, Biogen, Bristol-Myers Squibb, Covance, GE Healthcare, Genentech, GlaxoSmithKline, Lilly, Lundbeek, Merck, Meso Scale Discovery, Pfizer, Piramal, Roche, Servier, Teva, UCB, and Golub Capital.

## Notes

**Financial disclosures:** None

**Conflicts of interest**: On behalf of all authors, the corresponding author states that there is no conflict of interest.

